# The Impact of African and Brazilian Zika virus isolates on neuroprogenitors

**DOI:** 10.1101/046599

**Authors:** Loraine Campanati, Luiza M. Higa, Rodrigo Delvecchio, Paula Pezzuto, Ana Luiza Valadão, Fábio L. Monteiro, Grasiella M. Ventura, Carla Veríssimo, Ana M. Bispo De Filippis, Renato S. Aguiar, Amilcar Tanuri

**Author notes:** These authors contributed equally to this work (Loraine Campanati, Luiza M. Higa). Correspondence to: Amilcar Tanuri and/or Loraine Campanati.

## Abstract

In the last few months, an overwhelming number of people have been exposed to the Zika virus (ZIKV) in South and Central America. Here we showed, *in vitro*, that a Brazilian isolate impacts more severely murine neuronal progenitors and neurons than the African strain MR766. We found that the Brazilian isolate more pronouncedly inhibits neurite extension from neurospheres, alters their differentiation potential and causes neurons to have less and shorter processes. Comparing both lineages using a panel of inflammatory cytokines, we showed, with human neuroblastoma cells, that ZIKV induces the production of several inflammatory and chemotactic cytokines and once again, the Brazilian isolate had a more significant impact. Although much more needs to be studied regarding the association of ZIKV infection and brain damage during development, our study sheds some light into the differences between African and American lineages and the mechanisms by which the virus may be affecting neurogenesis.

## Main Text

In the last few months, healthcare professionals in Brazil noticed an alarming surge of fetuses diagnosed with microcephaly and several other brain malformations from mothers who showed typical signs of Zika virus infection during pregnancy (*1*). At the same time, retrospective statistical analysis from an outbreak (2013-2015) in the French Polynesia showed a correlation between Zika infection and microcephaly and also an increase in the number of patients with Guillain-Barré syndrome (*2, 3*).

Genetic and environmental agents have been linked to neurodevelopmental disorders, when they converge to defects on the mechanisms of proliferation, migration and differentiation of neural stem cells (NSCs) (*4, 5*). Congenital Cytomegalovirus infection is known to cause microcephaly by the inhibition of NSCs differentiation into neurons (*6*) and flaviviruses, like the Japanese encephalitis virus (JEV) infect neurons, glial cells and neural stem cells, leading to changes in their differentiation potential (*7-10*). Evidence is mounting on the correlation of ZIKV infection to damage to the nervous system, but there is a lack of information about the differences between African and Brazilian isolates on neural cells (*11, 12*). ZIKV particles were recently identified in the brain tissue of fetuses diagnosed with microcephaly and other brain damages, whose mothers were infected in the Americas (*12, 13*) and the MR766 African strain was shown to infect human neural stem cells derived from reprogrammed skin cells and to impair the growth of brain organoids (*14*) (P Garcez et al http://peerj.com/preprints/1817).

Sequencing and phylogenetic data from our group showed that isolates from Brazilian patients are 97-100% identical to the virus isolated from the outbreak in the French Polynesia, while showing 87-90% identity to African isolates (*11*). To date, there is no report of either cases of congenital abnormalities or Guillain-Barré syndrome in Africa, pointing to the possibility that differences in genome sequences of these strains may be associated to different clinical outcomes. In this work, we sought to comparatively investigate the outcomes of the infections of African and Brazilian ZIKV isolates in different cell types.

When the polyprotein sequence from prototypic African isolate MR766 was compared to the pandemic strain circulating in South America, we found an overall identity of 96.4 % and a 98.4% of similarity. Comparisons of the structural and non-structural proteins of the Brazilian isolate ZIKV Br 3189 (*11*) and MR766 showed 129 amino acids changes. Most of these substitutions are conservative and mainly located in the envelope and NS4b proteins, two critical points of the viral proteins. The first main modification acquired by ZIKV Br, is a new, canonical N-Linked glycosylation site located at position 153 of the envelope protein like recently described by cryo-electronmicroscopy (*15*). Another noticeable cluster of mutation is located at the cytoplasmic loop of the NS4b protein. This site is implicated in the induction of membrane synthesis and replication competence in Dengue virus (*16*), as well as in signal transduction related to malignant transformation in HCV (*17*) (Figs. S1-S2). These two main differences are found in all ZIKV isolates sequenced in the Americas and deposited in the Genbank site. These unique properties of the Asian lineage showed here could impact the field of vaccine and antiviral development to this new pathogenic agent.

Using both Brazilian and African isolates, we infected neurospheres derived from murine cortex, SH-SY5Y cells (human neuroblastoma cell line), mouse cortical neural stem cells and neurons.

We performed qPCR targeting ZIKV genome (supp. methods) and verified the intracellular presence of total viral RNA and negative-sense RNA strand, the template for viral genome replication, revealing that all cell types were permissive and productively infected by both viruses (Supp. table 1). MR766 infection of human NSCs was described recently (*14*) and here we showed that ZIKV Br replication in rodent NSCs was more effective than ZIKV MR766. In contrast, MR766 produced higher levels of intracellular RNA than ZIKV Br in neurospheres enriched in neuronal progenitors and SH-SY5Y cells, whose culture is known to have neuroblast-like cells (*18*).

Neurospheres were infected to up to 72h (Fig. 1) and we found that infection with both isolates decreased their area (Fig 1G): ZIKV Br by 21% (3099 ± 211 µm^2^), while MR766 had a less pronounced effect: 9% (3548 ± 203 µm^2^). By phase contrast and confocal microscopy, mock-infected neurospheres (3907 ± 239 µm^2^) showed exuberant thin cell processes (Fig. 1A) with positive staining for microtubule-associated protein 2 (Map2), a marker of neuronal differentiation (Fig. 1B). MR766 infected neurospheres also exhibited Map2^+^ neurites, (Figs. 1C, 1D) although we noticed that in small neurospheres projections were often misshapen and convoluted (Fig. S5). ZIKV Br infection greatly reduced the amount of Map2^+^ cellular processes and we noticed Map2^-^cells with progenitor-like morphology migrating out of the spheres (Fig. 1E, 1F and Fig S4).

**Fig. 1.**
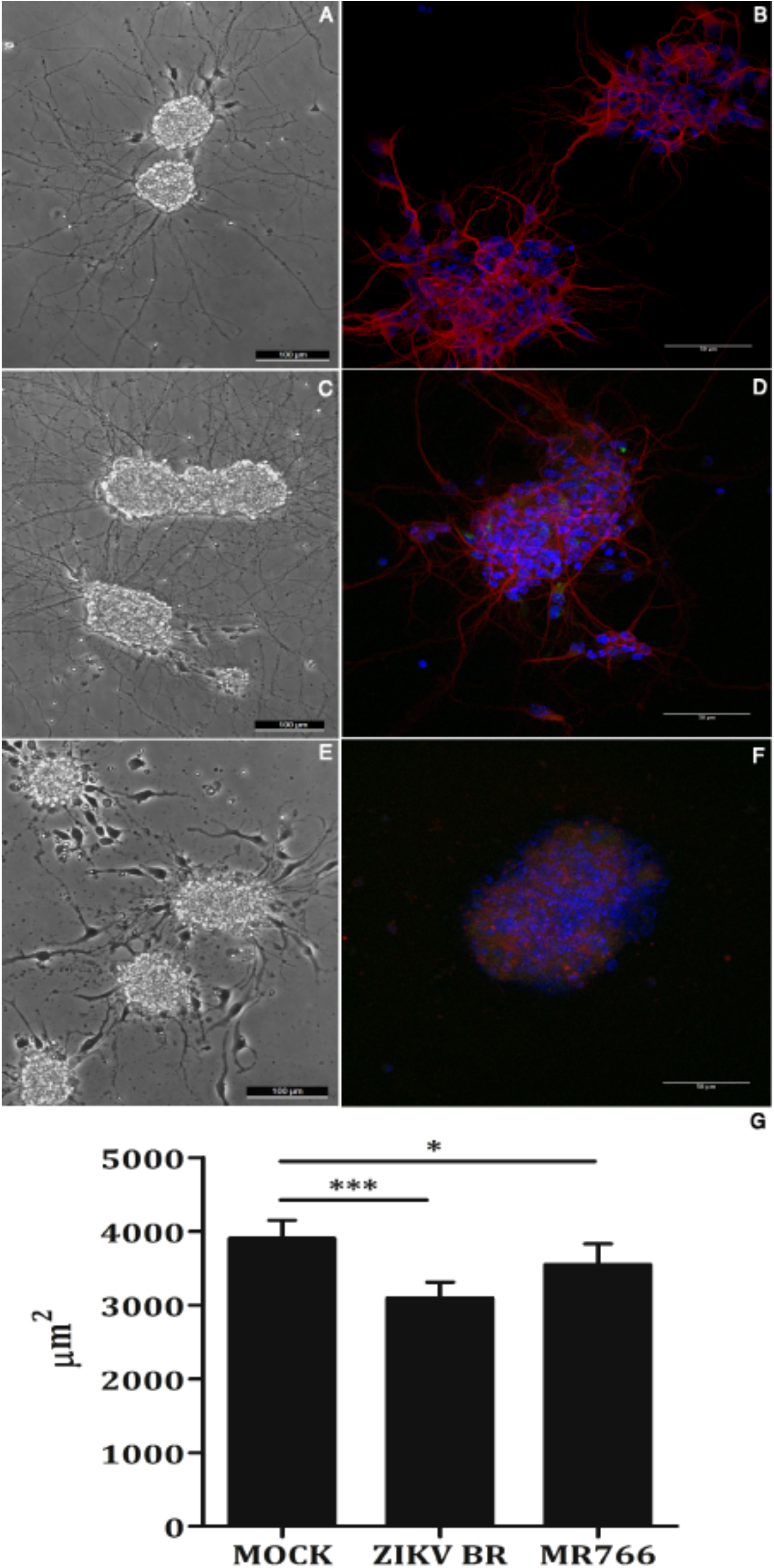
Infection with ZIKV Br inhibits Map2 process elongation and decreases the area of neurospheres. Mock infected neurospheres extend long, Map2^+^ processes (A,B), while MR766 infected suffered little with viral infection (C,D). Spheres infected with ZIKV Br (E,F) showed more drastic effect and only few neurites were seen. Mean area of the neurospheres in µm^2^ (g) ± s.e.m. ***p<0,001; *p<0,05; Student’s t test.

Differentiated cortical neurons were also infected (Fig. 2). Mock infected neurons (Fig. 2A) showed long, intricate neurites growing out of the cell body, while MR766 and ZIKV Br infected cells show less and simpler processes (Figs. 2B, 2C). No infection was observed in differentiated neuronal cells with both MR766 and ZIKV Br evidenced by IF using an anti-Flavivirus (4G2) antibody (Supp. methods), but rather the labeling of cells with shorter Map2+ processes (Figs. 2B, green). In neurospheres, using a higher viral load of MR766, we observed Map2low cells stained with 4G2 (Figs. 2D-E), mainly in the periphery of the spheres. Only rarely, we observed double-labeled (Map2/4G2) processes leaving the neurospheres (Fig. S3A and B), which is in accordance with data published recently showing that the ZIKV infection rate in human neurons is low (*14*).

**Fig. 2.**
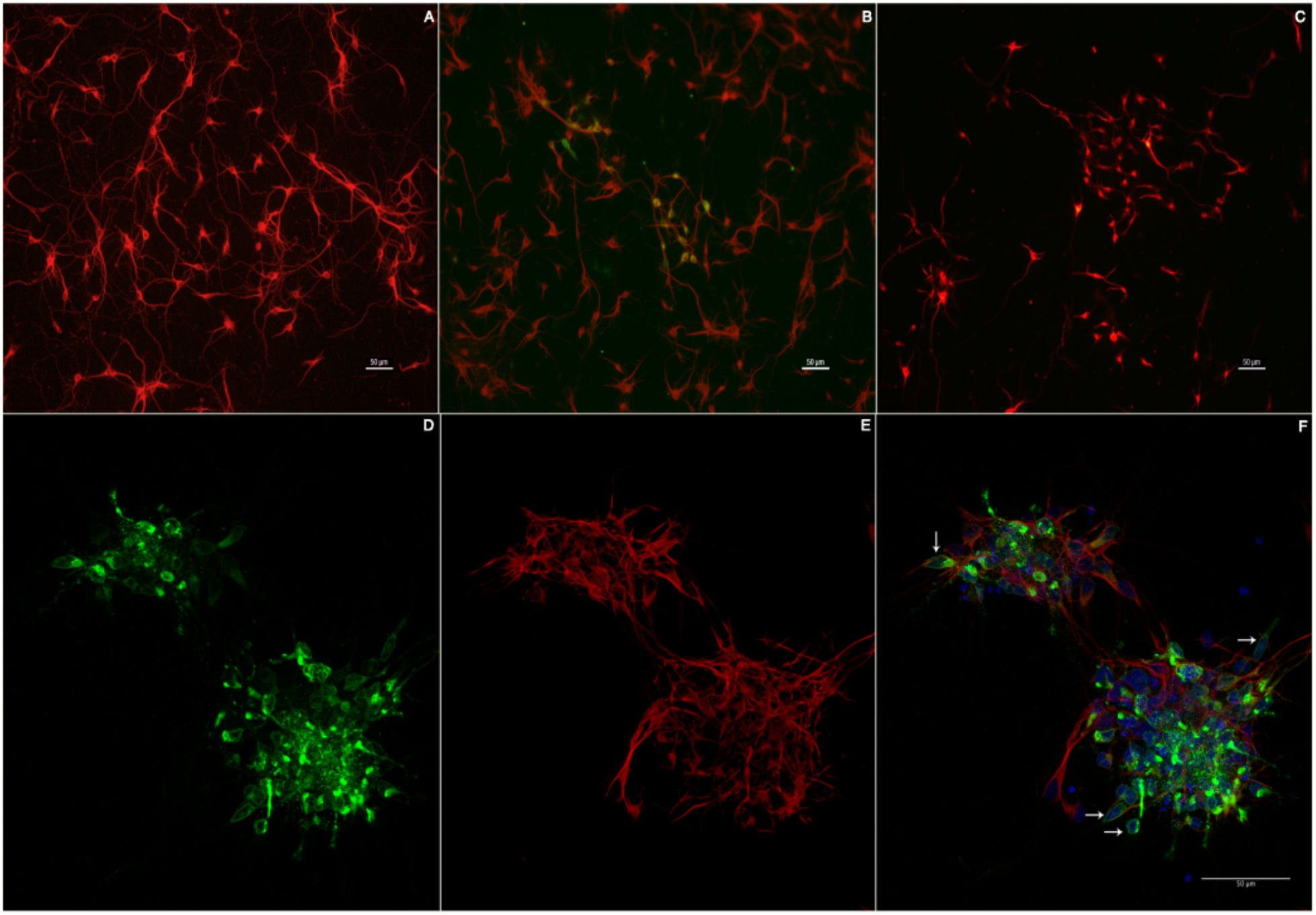
ZIKV infection decreases neuronal branching in culture. Neurons infected with both ZIKV isolates showed less branching (B, MR766; C, ZIKV Br) than Mock infected cells. ZIKV specific antigen labeling was detected specially in cells with few Map2+processes (B, green). At higher viral loads of MR766 (D-F), we noticed Map2low and Map2- cells (arrows) expressing viral antigens (green).

Taken together, these results show that both viruses are neurotropic, but have different replicative patterns depending on the differentiation status of the host cell, ZIKV Br being more replicative in undifferentiated cells than ZIKV MR766. Interestingly, even with lower levels of ZIKV Br replication in neurospheres, we observed more pronounced effects in neurite outgrowth by this isolate than when compared to ZIKV MR766. This result suggests a stronger impact in neuronal differentiation induced by the Brazilian isolate, which can be of great importance when one takes into account the outcome of infections in earlier phases of the nervous system development.

Other flaviviruses infections are associated to elevated levels of proinflammatory cytokines and chemokines that contribute to brain damage (*7*, *19*-*21*). To compare the immune activation caused by both viruses, we investigated the cytokine profile in the supernatant of SH-SY5Y cells infected with MR766 or ZIKV Br and found a significant increase of several cytokines when compared to mock infected cells (Supplementary Table 2). Two pro-inflammatory cytokines IL-1β and TNFα, four chemokines IL-8, IP-10, RANTES, Eotaxin and growth factors FGF2 and G-CSF were detected in higher concentration in the supernatant of ZIKV Br infected cells, when compared to cells infected with MR766. Unlike expected, most of the cytokines were not modulated by MR766 infection, except the pro-inflammatory IL-6 that was tightly down regulated (Fig. 3).

**Fig. 3.**
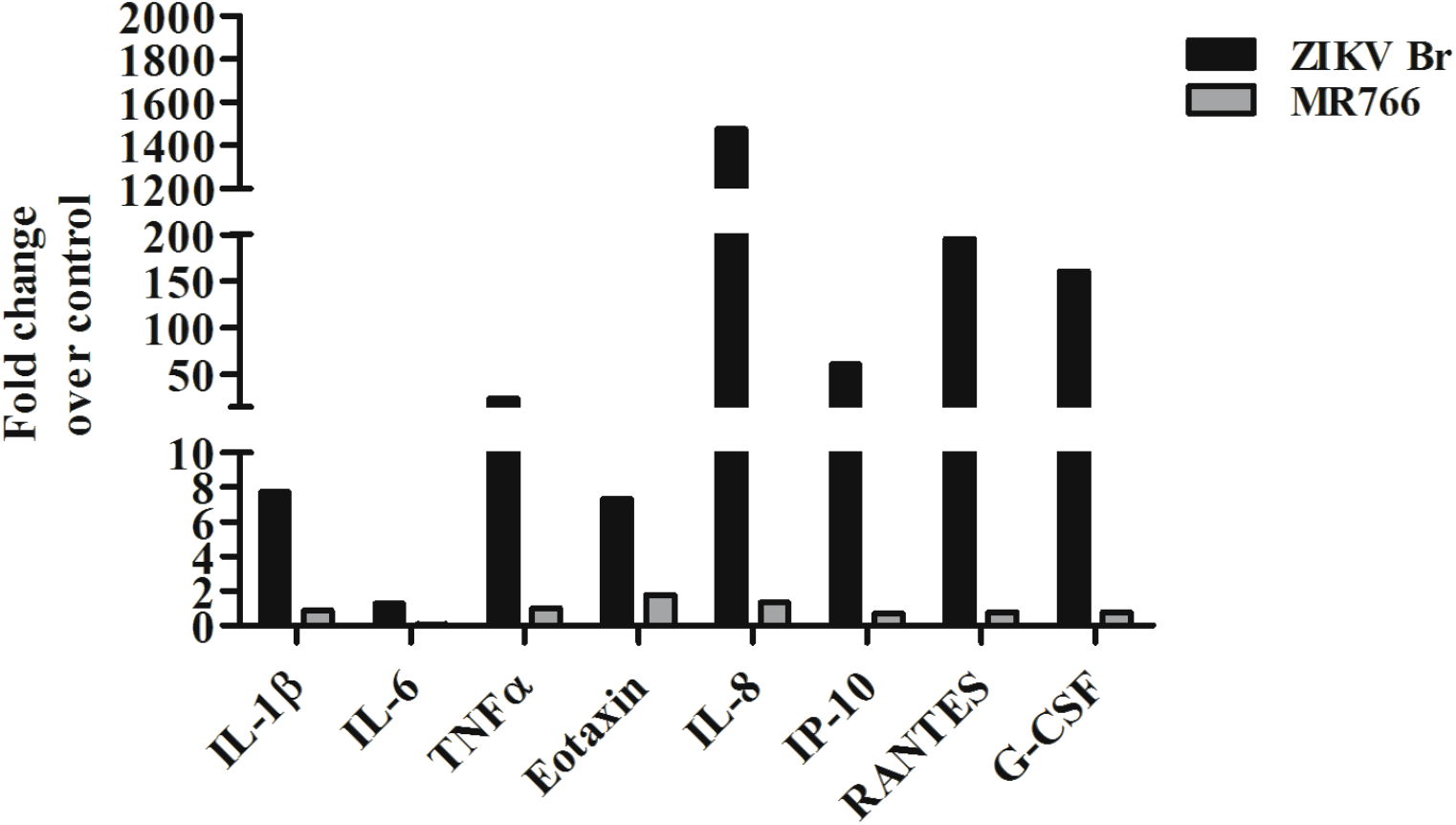
ZIKV Br infection elicits higher levels of cytokine production when compared to ZIKV 766 infection. A detailed table of all cytokines tested can be found in the supplementary methods.

IL-1β and TNFα act as potent mediators of neuronal death in several neurological disorders, West Nile virus and JEV infections (*9*). In neuroprogenitor cells it has been shown that IL-1β decreases cell proliferation while TNFa induces cell death via caspase-8-mediated apoptosis (*22*). Moreover, high levels of IL-8 have been shown to induce pro-apoptotic proteins and cell death in cultured neurons (*23*). RANTES levels are associated with viral encephalites and can be induced by NF-kB activation via IL-1β and TNFa stimulus in glial cells (*20*, *24*). IP-10 is a chemokine induced by viral infection in astrocytes that can regulate IFNy levels (*25*). In our experiment, IFNy had a slight increase despite of IP-10 levels. Eotaxin high levels are also associated to neurodegenerative disorder (*26*). The growth factors FGF2 and G-CSF are important cytokines involved in NSC proliferation and neurogenesis providing neuroprotection and decreasing apoptosis (*27*, *28*). These results show that, when the infection by African and Brazilian isolates are compared, ZIKV Br is capable of elicit a stronger immune response, which can also be of significant importance in the pathogenesis of the infection. Cytokine profile analysis can help us understand how immune response stimulated by ZIKV infection could disturb neuroprogenitor cells and contribute to the higher numbers of microcephaly and encephalitis cases described in South America and Asian countries with ZIKV outbreaks (*29*).

Overall, it is becoming increasingly clear that Zika virus is neurotropic as other flaviviruses. It is not known, however, whether the Brazilian isolate infectivity is different from the African strain and to which extent immune responses are involved in the pathogenesis. Work with animal models will certainly help to clarify theses questions. The work presented here indicates that the ZIKV circulating in South America, belonging to Asian lineage, gained a novel capacity of altering neuroprogenitor cells when compared to the isolate from the Zika forest in Uganda in 1947.

## Acknowledgments

The author would like to thank Conselho Nacional de Desenvolvimento e Pesquisa (CNPq), Fundação de Amparo a Pesquisa do Estado do Rio de Janeiro (FAPERJ) e Departamento de DST, AIDS e Hepatites Virais do Ministério da Saúde do Brasil for funding research in our laboratories.

---

**Figure S1.**
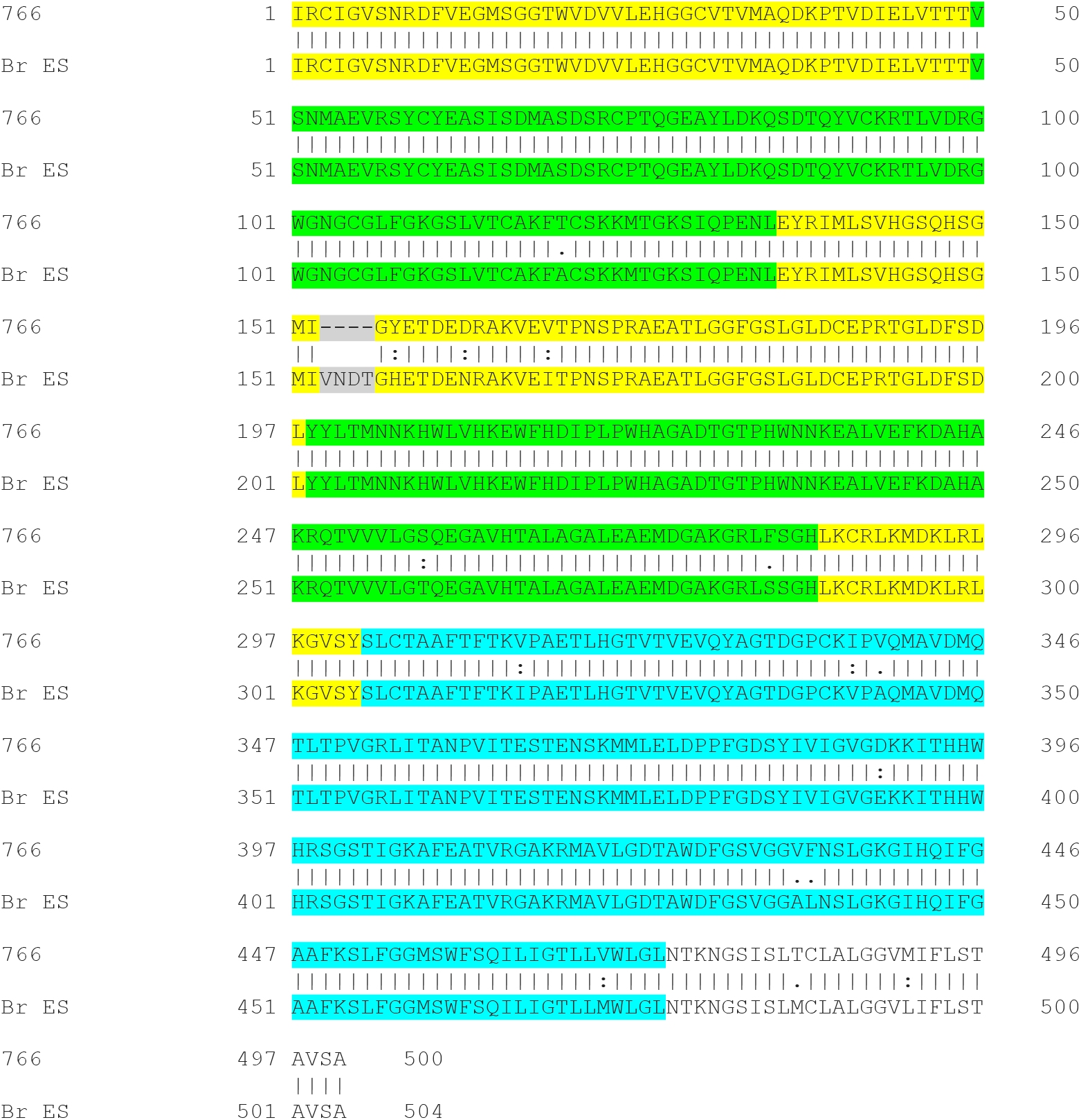
Amino acid alignment of ZIKV envelope sequence from African 766 (766) prototypic sequence with Brazilian (Br ES) counterpart done with *Needle* package contained in Clustal W package http://www.ebi.ac.uk/Tools/psa/emboss_needle/). Marked in yellow, green and blue are protein Domains I, II, and III, respectively. The insertion of four amino acid in BR ES Domain I is marked in gray. A vertical line between sequences represents identity and two and one points represents conservative and non-conservative substitutions, respectively. Total identity was 96.2% and similarity score was 98%.

**Figure S2.**
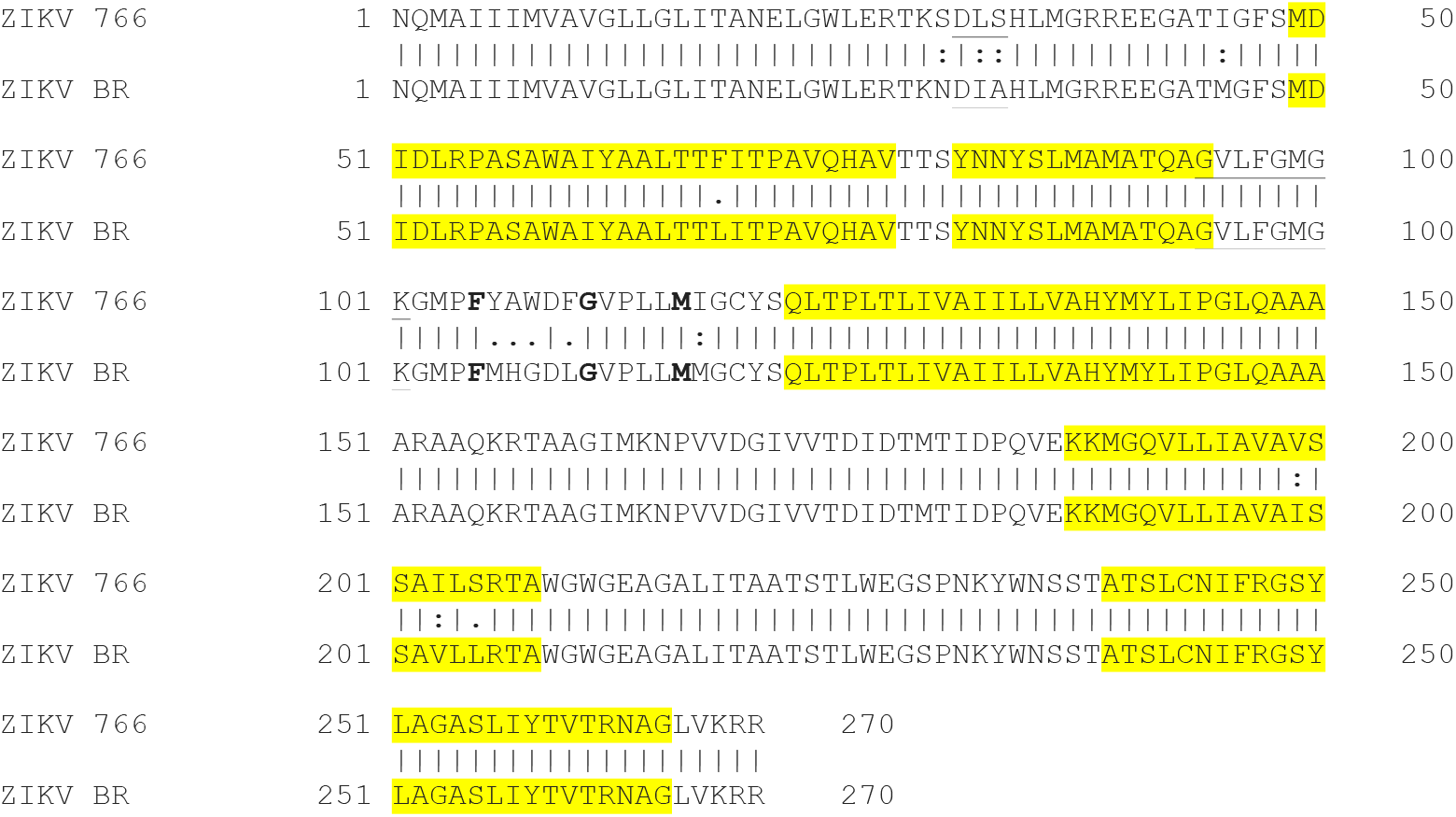
Amino acid alignment of ZIKV NS4B sequence from African MRC 766 prototypic sequence with Brazilian (Br ES) counterpart done with *Needle* package contained in Clustal W package http://www.ebi.ac.uk/Tools/psa/emboss_needle/). Marked in yellow are five NS4B transmembrane domains. The underlined sequences represents two DNA binding sequences motifs and in bold are three important amino acids for Dengue virus replication. A vertical line between sequences represents identity and two and one points represents conservative and non-conservative substitutions, respectively. Total identity was 96.2% and similarity score was 98%.

**Figure S3.**
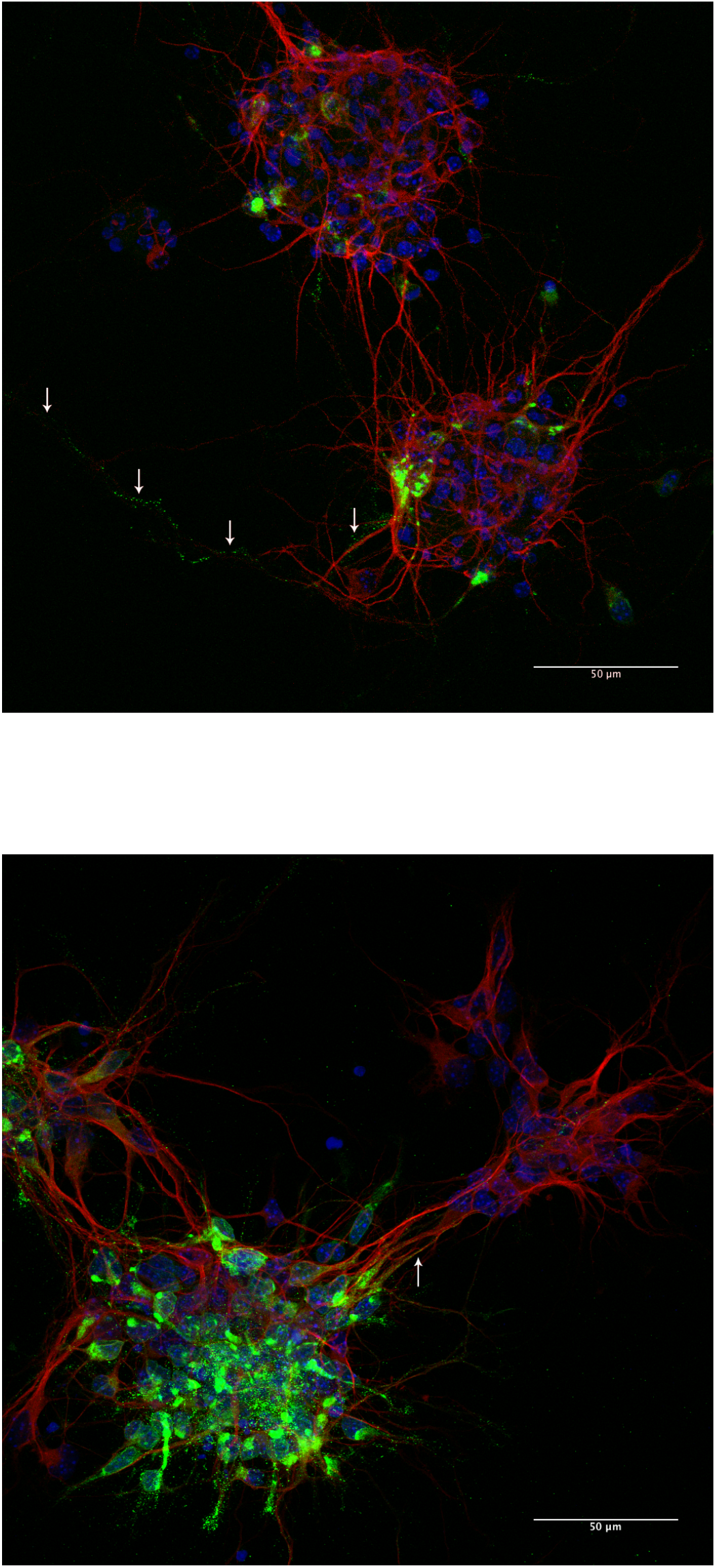
Neurospheres infected with 10^6^ p.f.u of ZIKV MR766 stained for Map2 (red) and flavivirus specific antigen (green). Only rarely we observed 4G2 positive neurites (arrows) growing out of the spheres.

**Figure S4.**
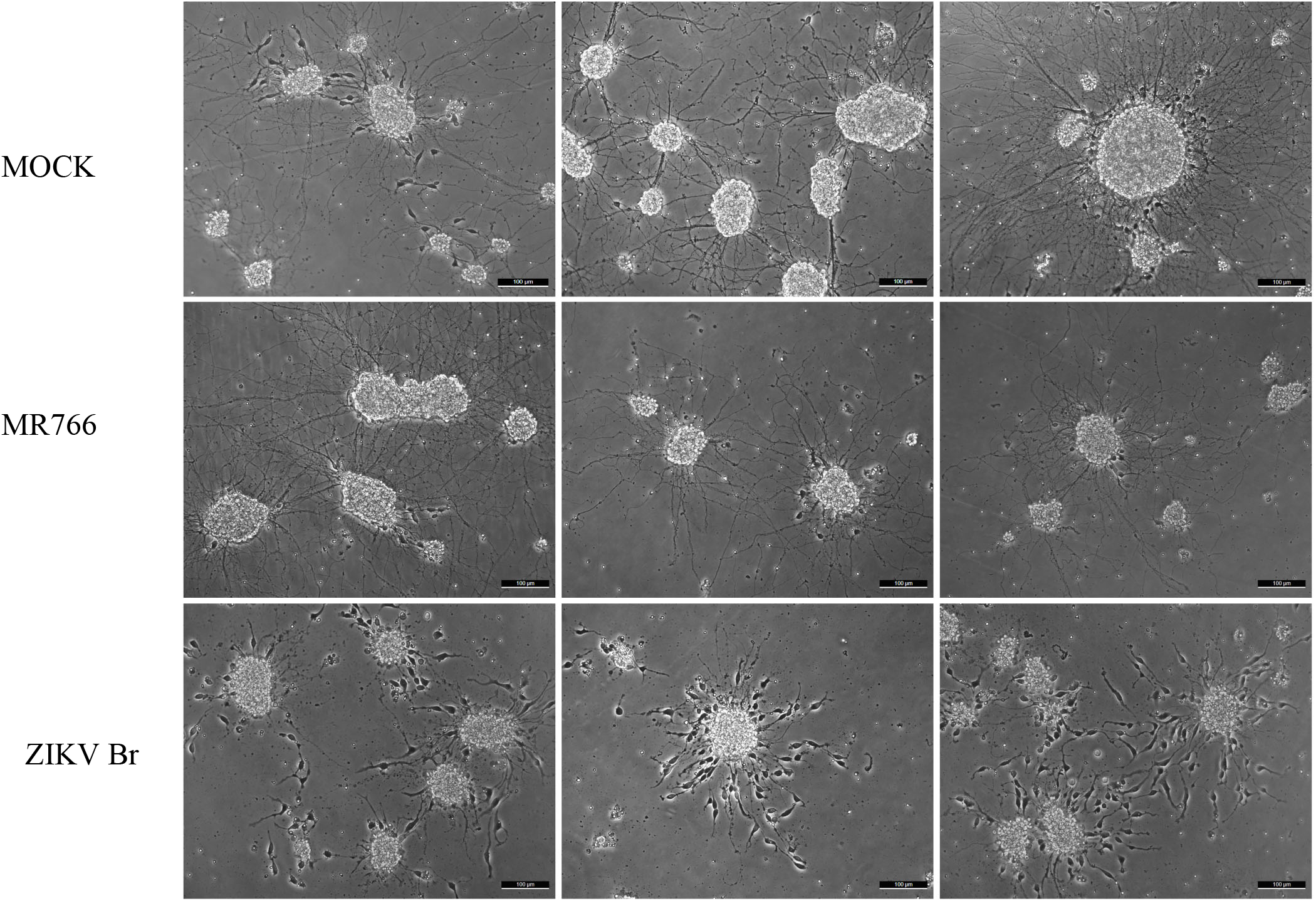
More examples of infected neurospheres as seen by phase contrast microscopy.

**Figure S5.**
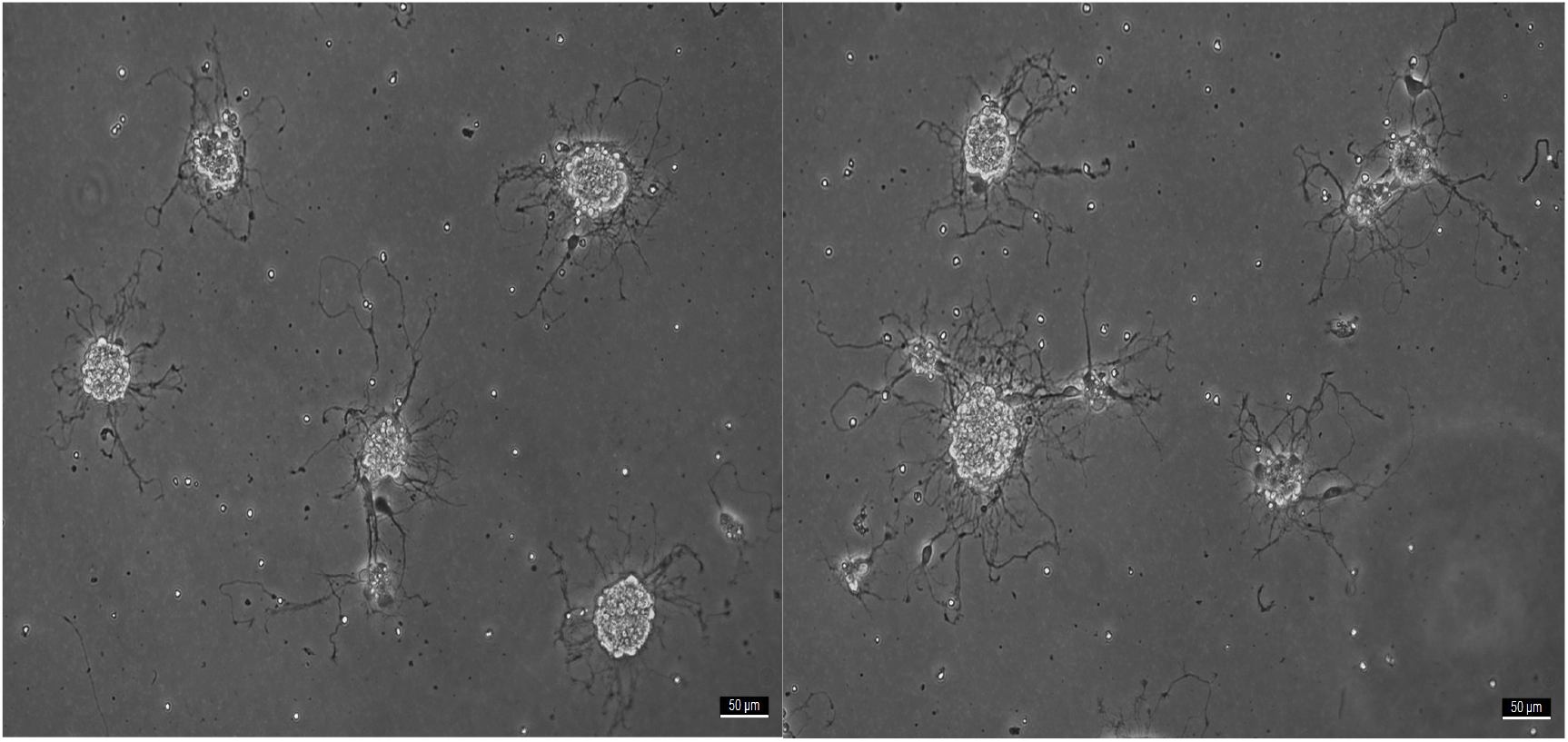
Smaller neurospheres infected with MR766 had misshapen and convoluted neurite outgrowth.

**Table S1.** Quantitative PCR of negative strand and total viral RNA from NSCs, SH-SY5Y and Neurosphere.

| Experiment condition | NSCs□ |  | SH-SY5Y□ |  | Neurosphere* |  |
| --- | --- | --- | --- | --- | --- | --- |
|  | RNA negative strand | Total RNA | RNA negative strand | Total RNA | RNA negative strand | Total RNA |
| Mock | - | - | - | - | - | - |
| ZIKV MR766 | + | + | +++ | +++ | ++ | ++ |
| ZIKV Br | ++ | ++ | ++ | +++ | + | + |

+ 30<CT<40; ++ 20<CT<30; +++ CT<20
□ virus infection with MOI 0.1
\*virus infection with 5x10<sup>4</sup> PFU

**Table S2.**
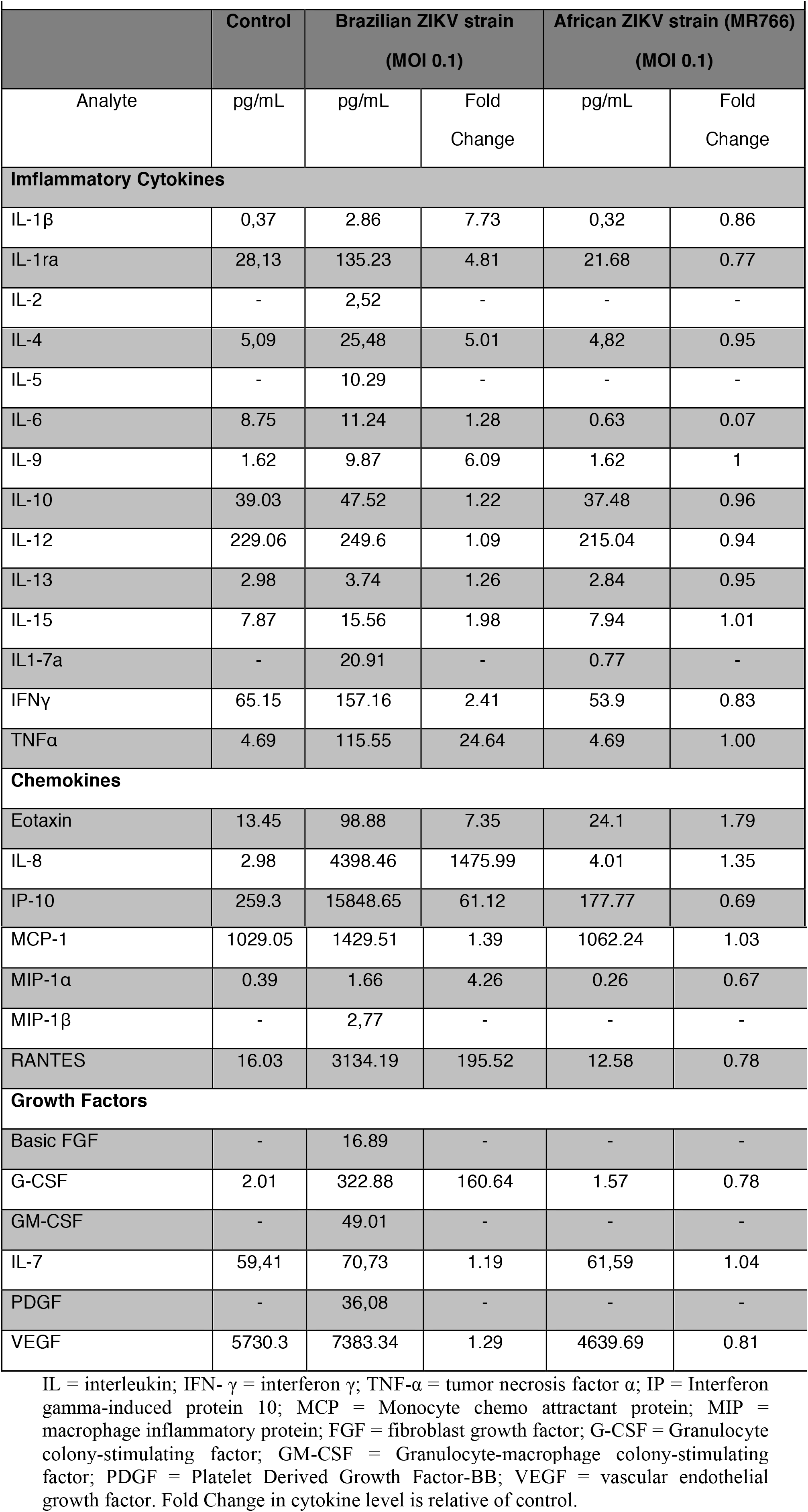
Comparison of Inflammatory Cytokines, Chemokines, and Growth Factors levels in the supernatant SH-SY5Y cells (neuroblastoma cell line) after infection with Brazilian and African ZIKV lineages.

